## supplemental for "Preserved reptile scales retain microscopic features, revealing a new instance of convergent evolution"

### S1 Supplementary Material

**Table S1:** Sample information. *Preserved in house by Joseph R. Mendelson III

| Species | Sample type | Year collected | Source |
|---|---|---|---|
| *Pantherophis guttatus* | shed skin (s1) | 2021 | Zoo Atlanta, Atlanta GA |
| *Pantherophis guttatus* | shed skin (s2) | 2024 | Rieser Lab, Emory University, Atlanta GA |
| *Pantherophis guttatus* | museum (m1) | unknown | University of Texas at Arlington, Arlington TX |
| *Pantherophis guttatus* | museum (m2) | 1964 | University of Texas at Arlington, Arlington TX |
| *Crotalus polystictus* | shed skin | 2024 | Zoo Atlanta, Atlanta GA |
| *Crotalus polystictus* | museum | 1939 | Field Museum of Natural History, Chicago IL |
| *Crotalus atrox* | shed skin (s1) | unknown | Courtesy of Gordon Schuett |
| *Crotalus atrox* | shed skin (s2) | unknown | Courtesy of Gordon Schuett |
| *Crotalus atrox* | museum* (p1) | 2021 | Courtesy of Gordon Schuett |
| *Crotalus atrox* | museum* (p2) | 2021 | Courtesy of Gordon Schuett |
| *Crotalus atrox* | frozen (f1) | 2021 | Courtesy of Gordon Schuett |
| *Crotalus atrox* | frozen (f2) | 2021 | Courtesy of Gordon Schuett |
| *Thamnophis saurita* | shed skin | unknown | Joseph R. Mendelson III |
| *Thamnophis saurita* | museum | 1971 | University of Texas at Arlington, Arlington TX |
| *Ophisaurus attenuatus* | shed skin | 2022 | Rieser Lab, Emory University, Atlanta GA |
| *Ophisaurus attenuatus* | museum | 1994 | University of Texas at Arlington, Arlington TX |
| *Cerastes cerastes* | shed skin | unknown | Joseph R. Mendelson III |
| *Cerastes vipera* | shed skin | unknown | Joseph R. Mendelson III |
| *Crotalus cerastes* | shed skin | unknown | Zoo Atlanta, Atlanta GA |
| *Bitis peringueyi* | museum | 1987 | University of Texas at Arlington, Arlington TX |

**Table S2:** JS divergence values between each pair of height distributions analyzed for each *P. guttatus* image. Note that JS divergence values between museum samples and shed skin samples are low and comparable to the values between shed skins indicating a high degree of similarity.

| samples | **s1a** | **s1b** | **s1c** | **s1d** | **s1e** | **s1f** | **s2** | **m1** | **m2** |
|---|---|---|---|---|---|---|---|---|---|
| **s1a** | 0 | 0.0273 | 0.0393 | 0.0157 | 0.0185 | 0.0104 | 0.0423 | 0.0332 | 0.0149 |
| **s1b** | 0.0273 | 0 | 0.0021 | 0.0048 | 0.0018 | 0.0054 | 0.0076 | 0.0042 | 0.0184 |
| **s1c** | 0.0393 | 0.0021 | 0 | 0.0092 | 0.0054 | 0.0122 | 0.0069 | 0.0051 | 0.0277 |
| **s1d** | 0.0157 | 0.0048 | 0.0092 | 0 | 0.0012 | 0.0029 | 0.0193 | 0.0124 | 0.0186 |
| **s1e** | 0.0185 | 0.0018 | 0.0054 | 0.0012 | 0 | 0.0022 | 0.0125 | 0.0074 | 0.0155 |
| **s1f** | 0.0104 | 0.0054 | 0.0122 | 0.0029 | 0.0022 | 0 | 0.0162 | 0.0114 | 0.0095 |
| **s2** | 0.0423 | 0.0076 | 0.0069 | 0.0193 | 0.0125 | 0.0162 | 0 | 0.0039 | 0.0203 |
| **m1** | 0.0332 | 0.0042 | 0.0051 | 0.0124 | 0.0074 | 0.0114 | 0.0039 | 0 | 0.0175 |
| **m2** | 0.0149 | 0.0184 | 0.0277 | 0.0186 | 0.0155 | 0.0095 | 0.0203 | 0.0175 | 0 |

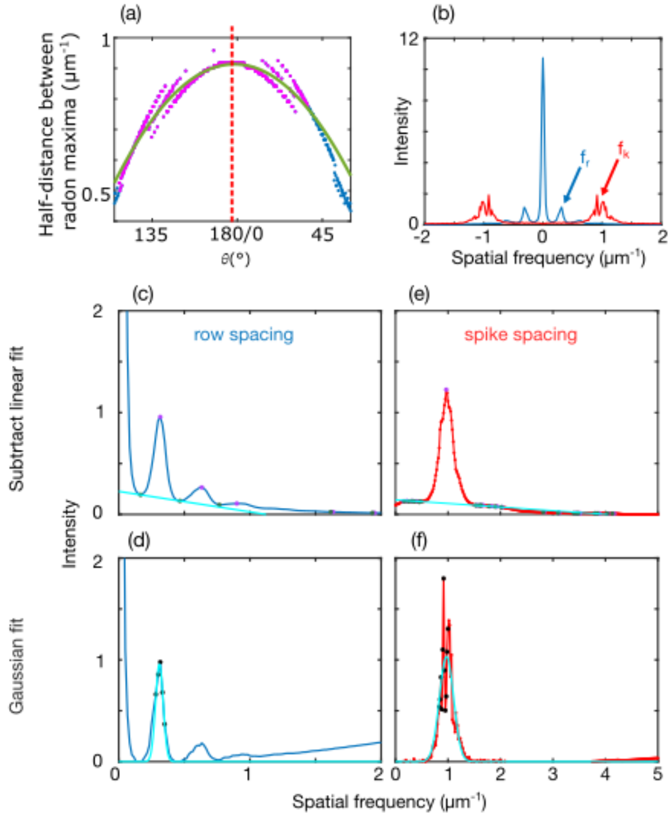


**Figure S1: Quantification of microscopic features shown in Figure 3 (a)** The distance between bands in each slice of the Radon transform are used to find the slice that gives the spatial frequency of spikes. At each angle value in the Radon transform, the half-distance between maxima on either side of zero was recorded and plotted. 400 points (magenta) around the maximum distance are fit with a parabola to account for noise. The angle value at the peak of the parabola is used to get the slice that gives the spike spacing. **(b)** Slices of the Radon transform used to calculate row spacing and spike spacing. The row spacing slice is the one that contains the maximum value in the whole Radon transform. **(c)** The peak of the Radon transform is set to a value of zero. To measure the row spacing, we take the positive half of the slice, smooth the data and find the first two local minima closest to zero. These minima surround a peak that gives the row frequently. We fit a linear model to these minima and subtract it from the data bringing the peak down to the x-axis to accurately assess its width. **(d)** We fit a Gaussian model to 5 points around the peak in the original, unsmoothed data. The peak position of the Gaussian is taken to be the row frequency and the standard deviation of the Gaussian is used as a measure of sample uncertainty. **(e)** A similar technique is applied to both sides of the spike spacing slice. Here a linear model is fit to the two minima surrounding the peak of the smoothed data and subtracted. **(f)** A Gaussian is fit to 11 points surrounding the peak in the unsmoothed data. After this method is repeated on the negative side of the slice, the spike frequency is taken to be the average absolute value of the two Gaussian peaks. The average standard deviation is used as a measure of sample uncertainty.

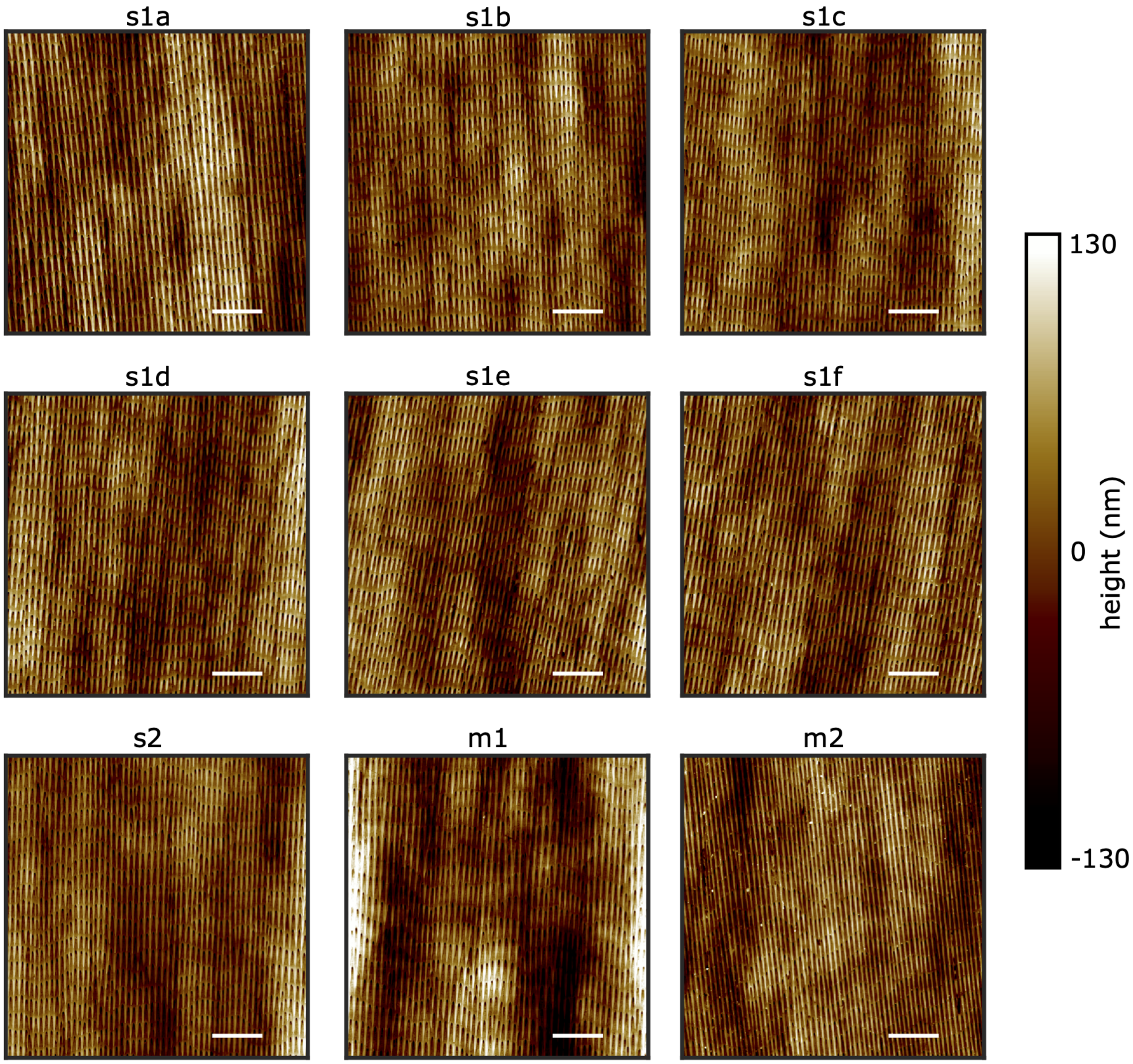


**Figure S2: Images from *P. guttatus* samples before a Gaussian filter is applied.** Images s1a-s1f are from six sites across the same shed skin sample. s2 shows a second shed skin sample. m1 and m2 show museum samples from two individuals. Images are used for analysis in Figures 3 and 4. Scale bars: 10$\mu$m

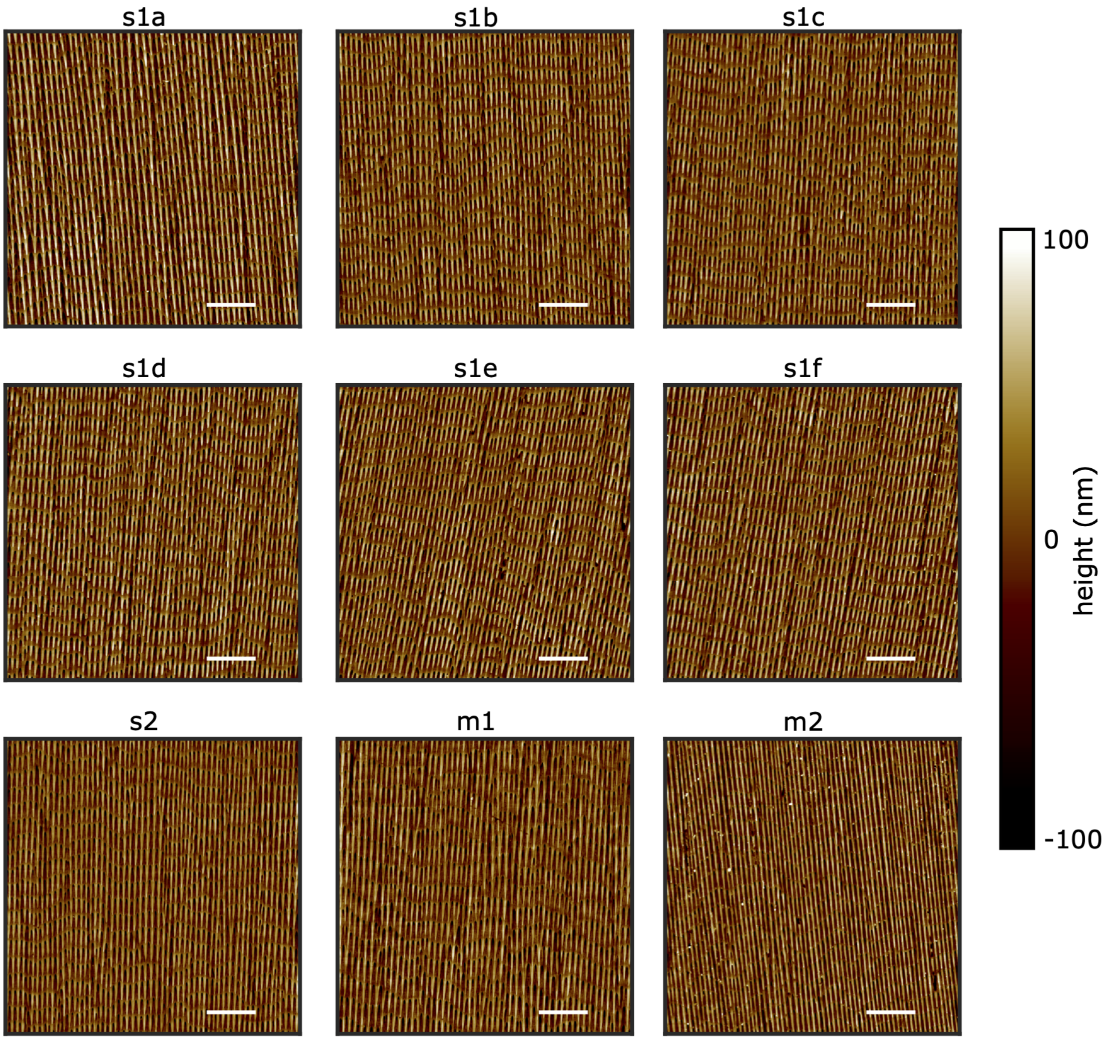


**Figure S3: Images from *P. guttatus* samples after a Gaussian filter is applied.** Images correspond to the labeled images in Figure S2. Images are used for analysis in Figures 3 and 4. Scale bars: 10$\mu$m

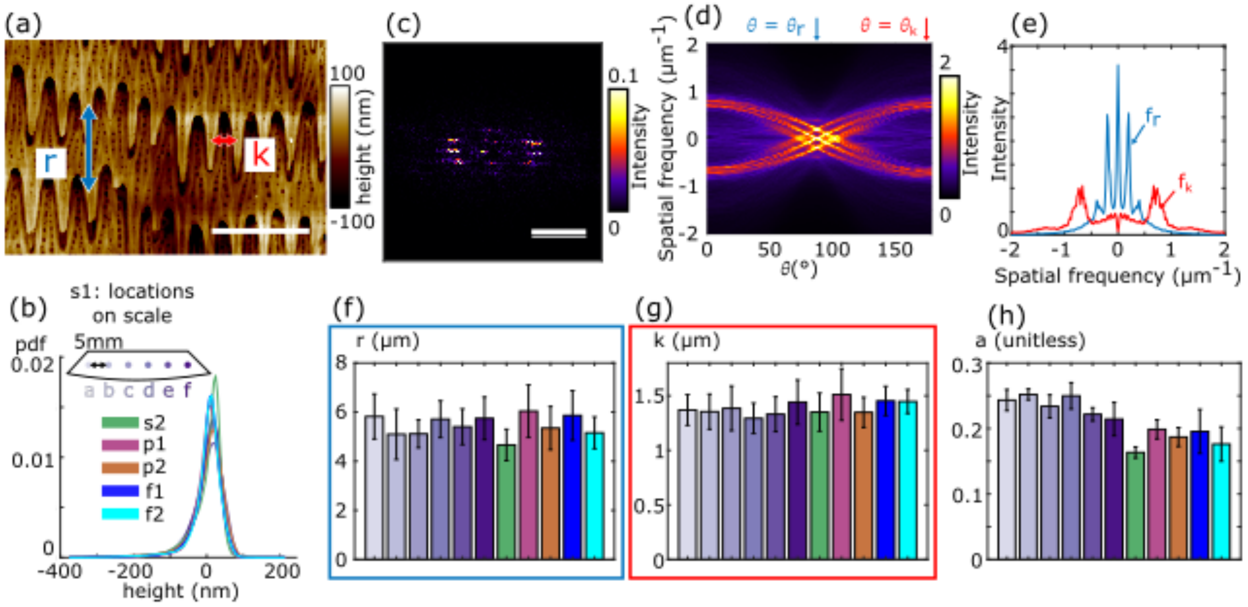


**Figure S4: The same analysis from Figures 3 and 4 with samples from *C. atrox*. This analysis also includes samples that are frozen and shows that microstructures are maintained on these as well.** **(a)** An image shows the details of the microstructure in sample s1. Height distributions from filtered $60\mu m \times 60\mu m$ images at 6 sites in s1 as well as single sites in s2, p1, p2, f1, and f2 **(b)** are similar. A power spectrum **(c)** and Radon transform **(d)** are calculated from each image and the row spacing and spike spacing are calculated from their corresponding Radon transform slices according to the method outlined in Figure S1. For this species, the row spacing **(f)** and spike spacing **(g)** are both retained in hand-preserved (p1, p2) as well as frozen (f1, f2) samples. The anisotropy **(h)**of each sample is slightly more variable but the anisotropy of the frozen and preserved samples are within the expected variation. Scale bars: **(a)** $2\mu m$, **(b)** $1\ \mu m^{-1}$

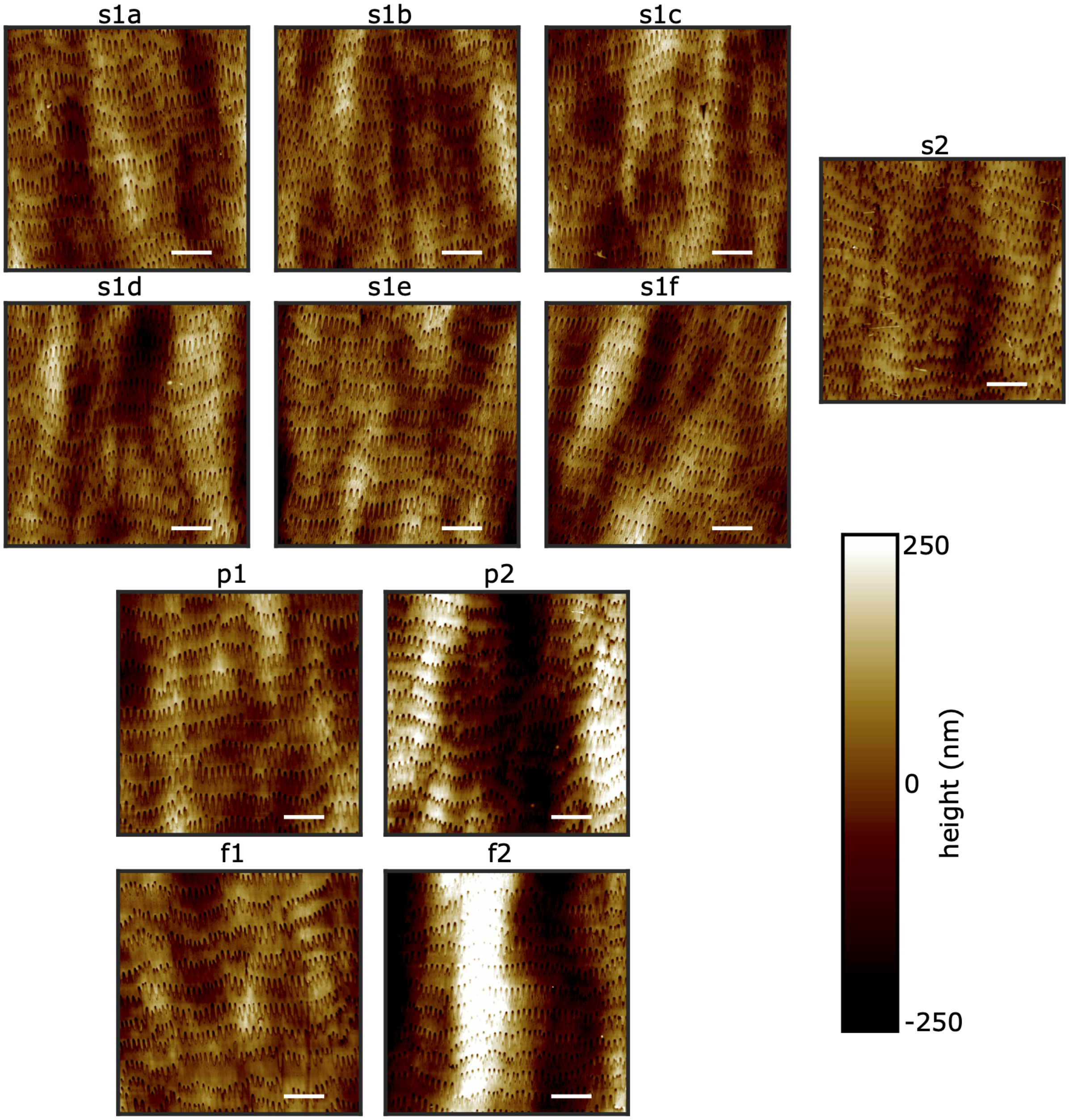


**Figure S5: Images from *C. atrox* samples before a Gaussian filter is applied.** Images s1a-s1f are from six sites across the same shed skin sample. s2 shows a second shed skin sample. p1 and p2 show samples that were preserved in-house using the same technique museum samples from two individuals. Samples f1 and f2 were frozen after mortality and thawed before imaging. Images are used for analysis in Figure S4. Scale bars: 10$\mu$m

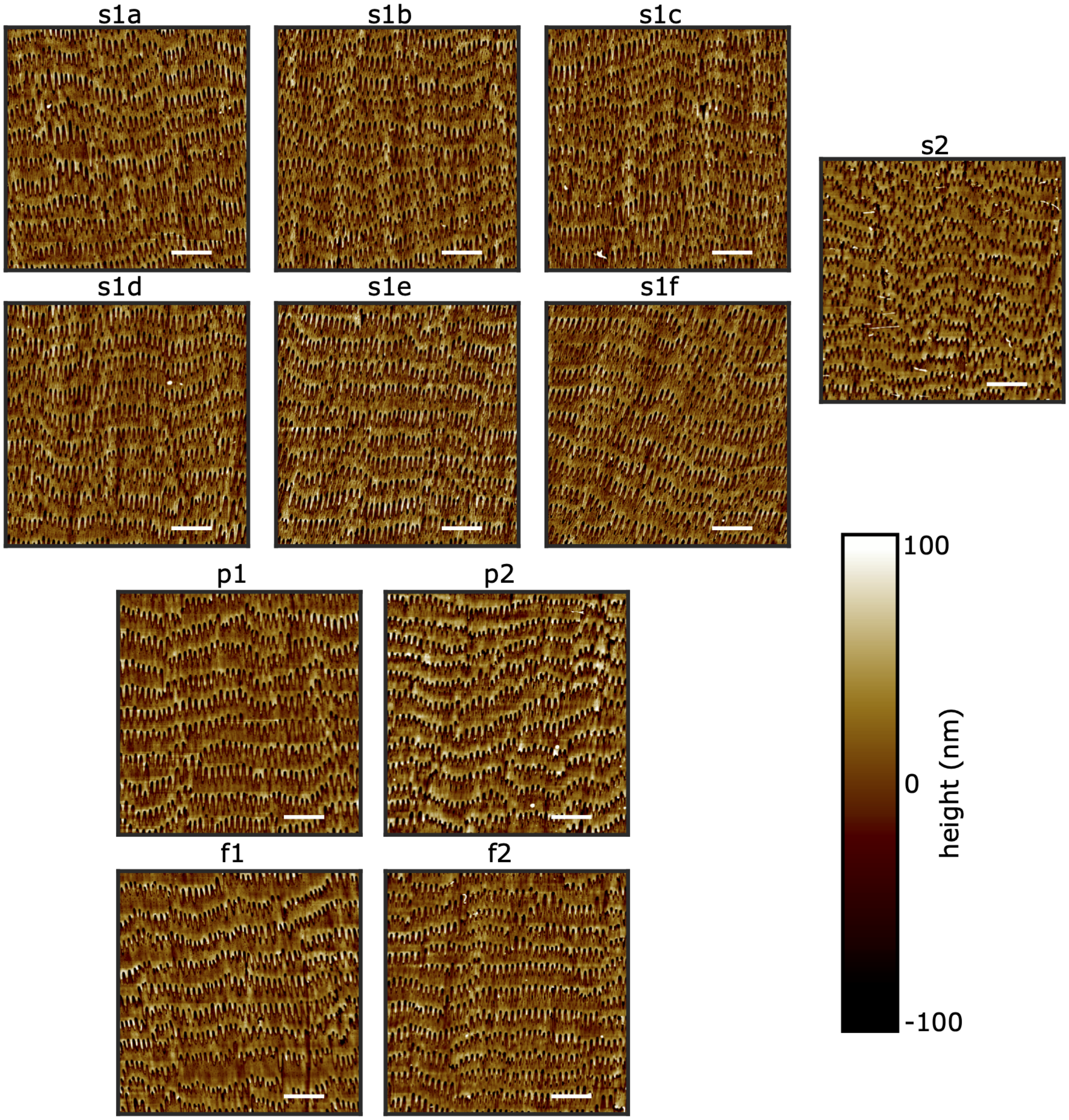


**Figure S6: Images from *C. atrox* samples after a Gaussian filter is applied.** Images correspond to the labeled images in Figure S5. Images are used for analysis in Figure S4. Scale bars: 10$\mu$m

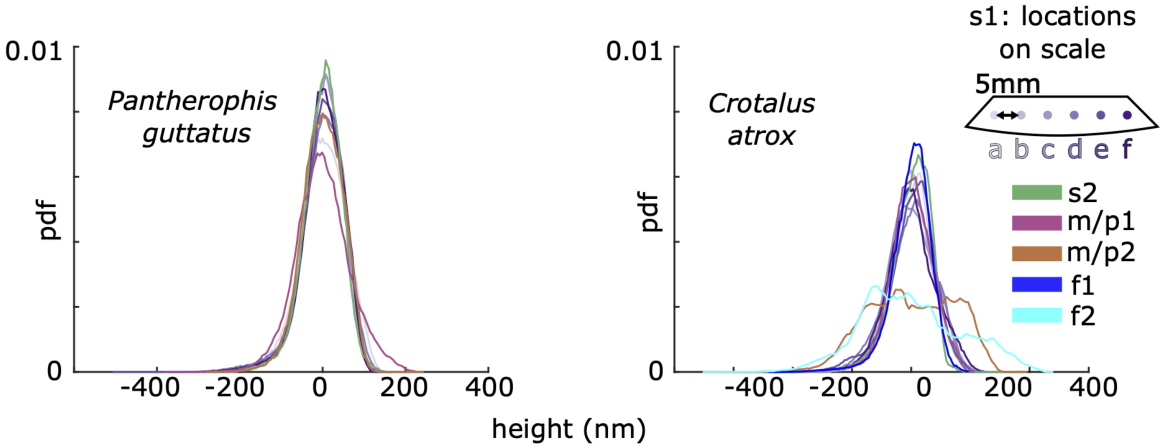


**Figure S7: Distributions of heights from unfiltered images.** Distributions from each image site in each sample from *P. guttatus* and *C. atrox* are shown.

**Table S3:** Spacing data for species where multiple sample types were studied. Most have spiked microstructures and strong periodicity in two directions. *O. attenuatus* only had strong periodicity in one direction corresponding to the rows of cells.

| species | sample/site label | row spacing $r$ | $\sigma_r$ | $\sigma_{rfit}$ | spike spacing $k$ | $\sigma_k$ | $\sigma_{kfit}$ | anisotropy | $\sigma_a$ |
|---|---|---|---|---|---|---|---|---|---|
| *P. guttatus* | s1a | 3.076 | 0.284 | 0.055 | 1.224 | 0.089 | 0.025 | 0.587 | 0.037 |
| *P. guttatus* | s1b | 3.249 | 0.303 | 0.045 | 1.063 | 0.058 | 0.011 | 0.535 | 0.019 |
| *P. guttatus* | s1c | 3.227 | 0.305 | 0.036 | 1.032 | 0.141 | 0.136 | 0.507 | 0.014 |
| *P. guttatus* | s1d | 3.140 | 0.288 | 0.049 | 1.006 | 0.043 | 0.004 | 0.540 | 0.016 |
| *P. guttatus* | s1e | 3.295 | 0.353 | 0.037 | 1.011 | 0.058 | 0.009 | 0.514 | 0.012 |
| *P. guttatus* | s1f | 3.222 | 0.406 | 0.054 | 1.111 | 0.084 | 0.013 | 0.542 | 0.011 |
| *P. guttatus* | s2 | 3.288 | 0.293 | 0.022 | 1.099 | 0.070 | 0.010 | 0.621 | 0.024 |
| *P. guttatus* | m1 | 4.101 | 0.443 | 0.038 | 1.228 | 0.110 | 0.033 | 0.595 | 0.014 |
| *P. guttatus* | m2 | 3.304 | 0.324 | 0.017 | 0.957 | 0.083 | 0.022 | 0.685 | 0.030 |
| *C. atrox* | s1a | 5.821 | 0.916 | 0.017 | 1.370 | 0.141 | 0.024 | 0.243 | 0.016 |
| *C. atrox* | s1b | 5.097 | 1.029 | 0.318 | 1.354 | 0.161 | 0.011 | 0.252 | 0.008 |
| *C. atrox* | s1c | 5.115 | 0.567 | 0.043 | 1.385 | 0.204 | 0.082 | 0.234 | 0.018 |
| *C. atrox* | s1d | 5.712 | 0.749 | 0.181 | 1.293 | 0.140 | 0.009 | 0.250 | 0.020 |
| *C. atrox* | s1e | 5.403 | 0.728 | 0.040 | 1.331 | 0.160 | 0.023 | 0.222 | 0.010 |
| *C. atrox* | s1f | 5.746 | 0.867 | 0.167 | 1.441 | 0.204 | 0.018 | 0.214 | 0.025 |
| *C. atrox* | s2 | 4.664 | 0.640 | 0.102 | 1.350 | 0.175 | 0.043 | 0.163 | 0.009 |
| *C. atrox* | p1 | 6.040 | 1.073 | 0.446 | 1.511 | 0.237 | 0.023 | 0.199 | 0.015 |
| *C. atrox* | p2 | 5.348 | 0.879 | 0.242 | 1.348 | 0.137 | 0.005 | 0.187 | 0.014 |
| *C. atrox* | f1 | 5.865 | 1.011 | 0.312 | 1.453 | 0.134 | 0.013 | 0.196 | 0.034 |
| *C. atrox* | f2 | 5.158 | 0.648 | 0.111 | 1.447 | 0.109 | 0.017 | 0.176 | 0.026 |
| *C. polystictus* | shed | 5.225 | 0.683 | 0.070 | 1.376 | 0.129 | 0.018 | 0.405 | 0.019 |
| *C. polystictus* | museum | 6.106 | 0.865 | 0.064 | 1.319 | 0.103 | 0.008 | 0.336 | 0.006 |
| *T. saurita* | shed | 4.273 | 0.542 | 0.035 | 4.304 | 0.743 | 0.078 | 0.614 | 0.024 |
| *T. saurita* | museum | 3.749 | 0.465 | 0.045 | 4.008 | 0.167 | 0.018 | 0.705 | 0.012 |
| *O. attenuatus* | shed | 3.640 | 1.269 | 0.098 | n/a | n/a | n/a | 0.103 | 0.040 |
| *O.attenuatus* | museum | 3.773 | 1.751 | 0.143 | n/a | n/a | n/a | 0.152 | 0.011 |

**Table S4:** Spacing data for sidewinder samples.

| species | sample type | pit spacing | $\sigma$ | $\sigma_{fit}$ | anisotropy | $\sigma$ |
|---|---|---|---|---|---|---|
| *Cerastes cerastes* | shed | 0.870 | 0.272 | 0.009 | 0.013 | 0.001 |
| *Cerastes vipera* | shed | 1.509 | 0.530 | 0.022 | 0.009 | 0.006 |
| *Crotalus cerastes* | shed | 0.843 | 0.339 | 0.009 | 0.010 | 0.004 |
| *Bitis peringueyi* | museum | 0.880 | 0.338 | 0.016 | 0.008 | 0.002 |

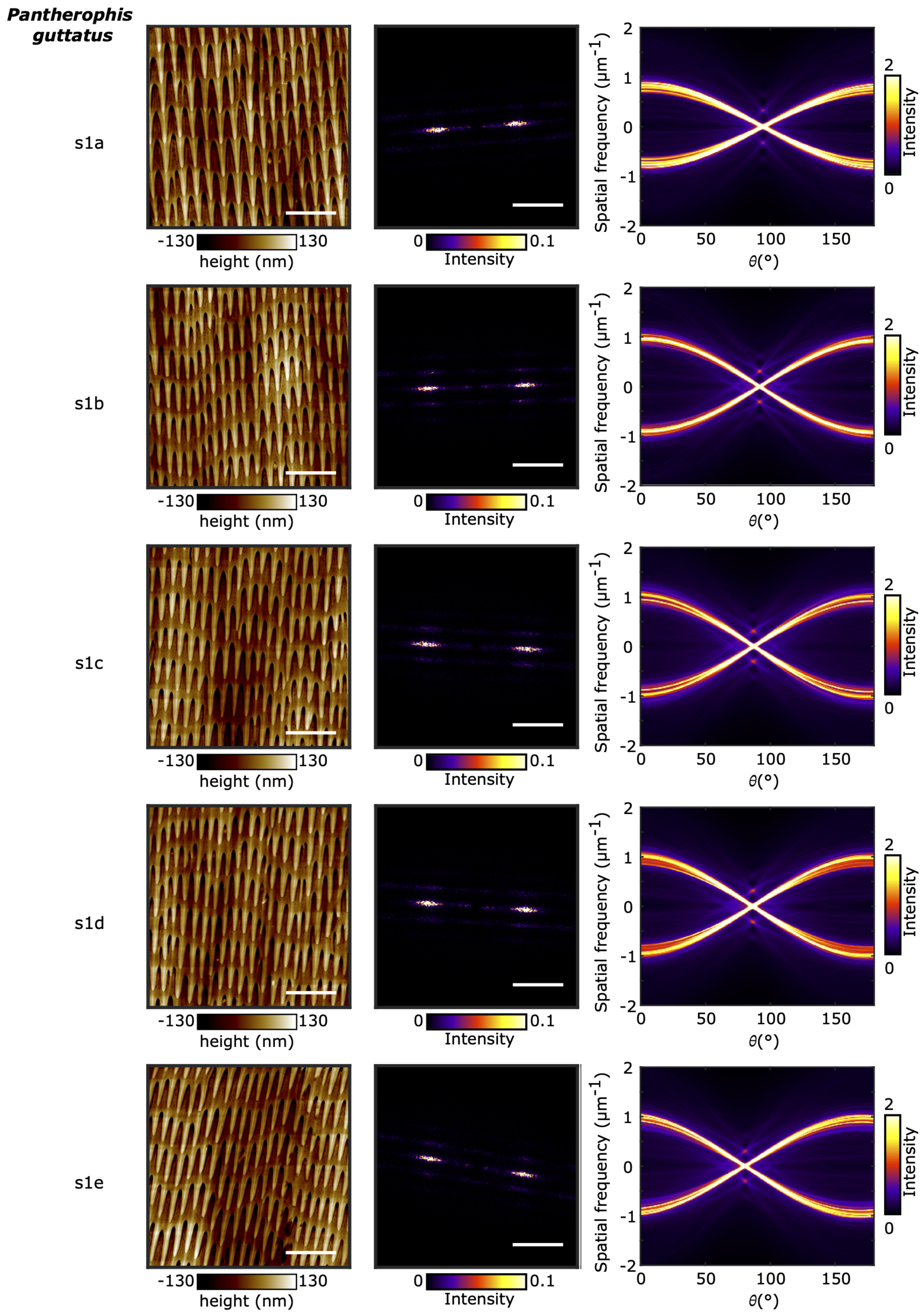


**Figure S8:** Images (20$\mu$m $\times$20$\mu$m), power spectra, and Radon transforms from *P. guttatus* samples. Scale bars: AFM image (left column): 5$\mu$m, power spectrum (middle column): 1$\mu$m$^{-1}$

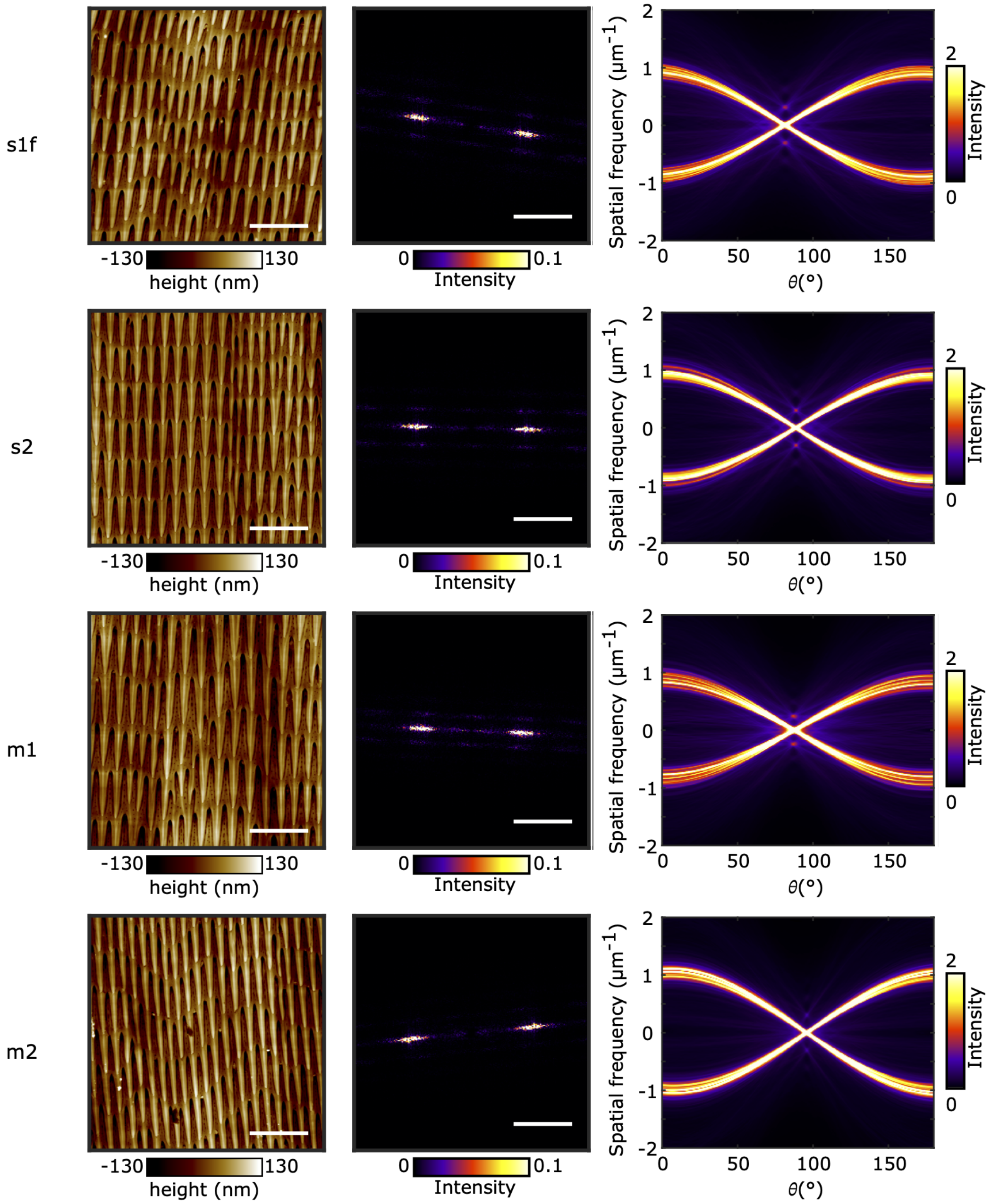


**Figure S9:** Cont. from Figure S8: Images (20$\mu$m $\times$20$\mu$m), power spectra, and Radon transforms from *P. guttatus* samples. Scale bars: AFM image (left column): 5$\mu$m, power spectrum (middle column): 1$\mu$m$^{-1}$

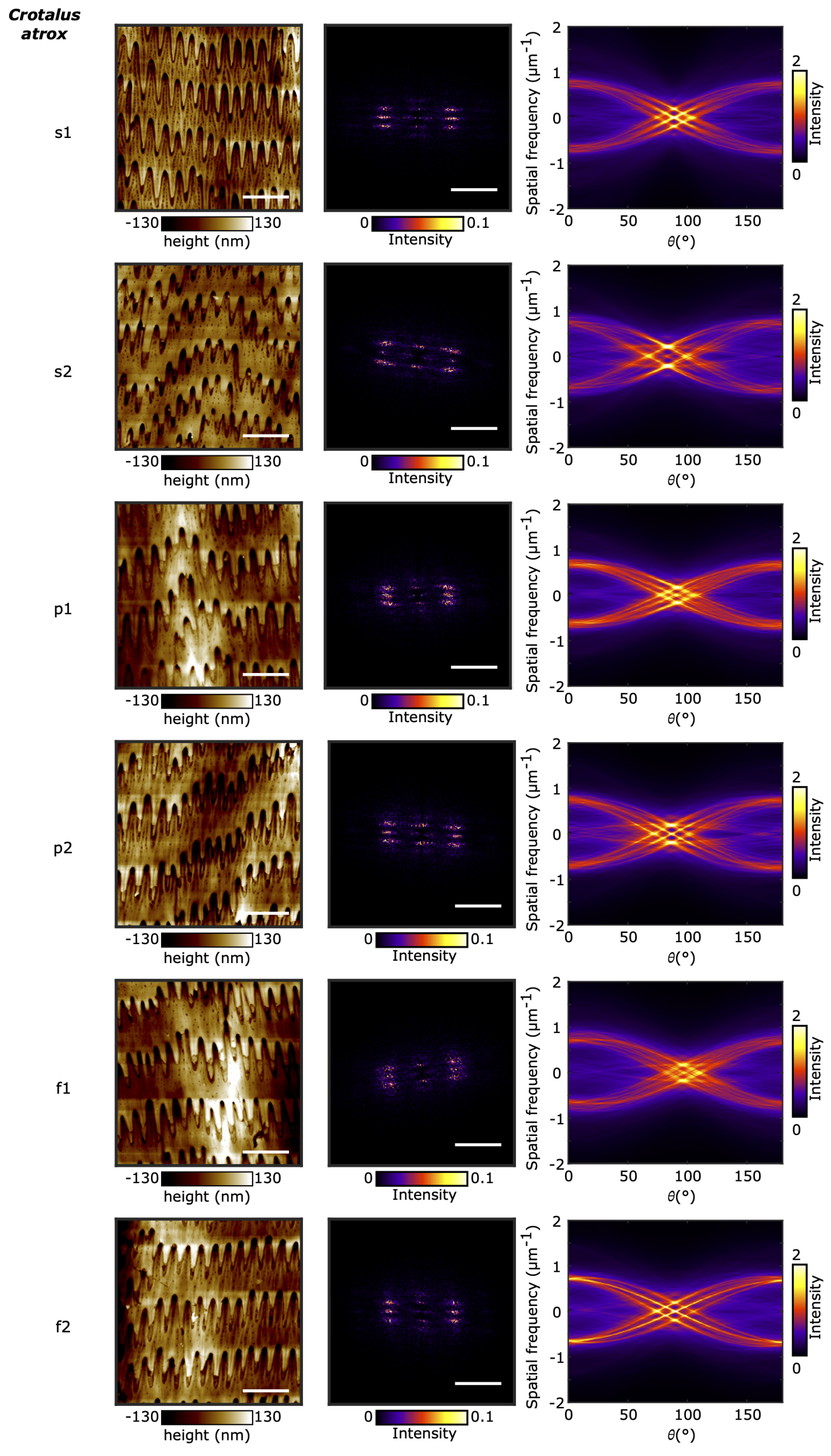


**Figure S10:** Images ($20\mu m \times 20\mu m$), power spectra, and Radon transforms from *C. atrox* samples. Scale bars: AFM image (left column): $5\mu m$, power spectrum (middle column): $1\mu m^{-1}$

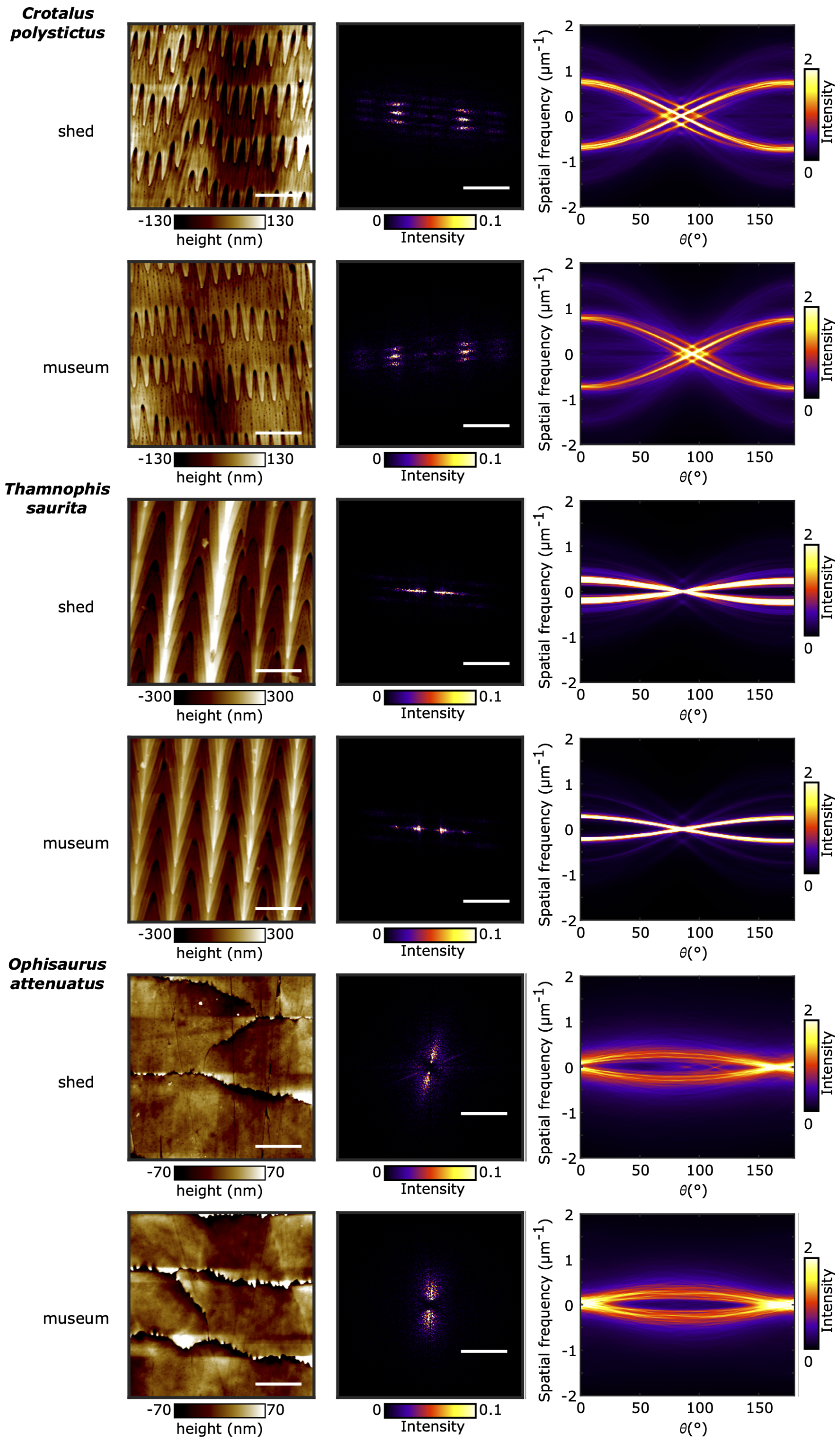


**Figure S11:** Images (20$\mu$m $\times$20$\mu$m), power spectra, and Radon transforms from *C. polystictus*, *T. saurita*, and *O. attenuatus* samples. Scale bars: AFM image (left column): 5$\mu$m, power spectrum (middle column): 1$\mu$m$^{-1}$

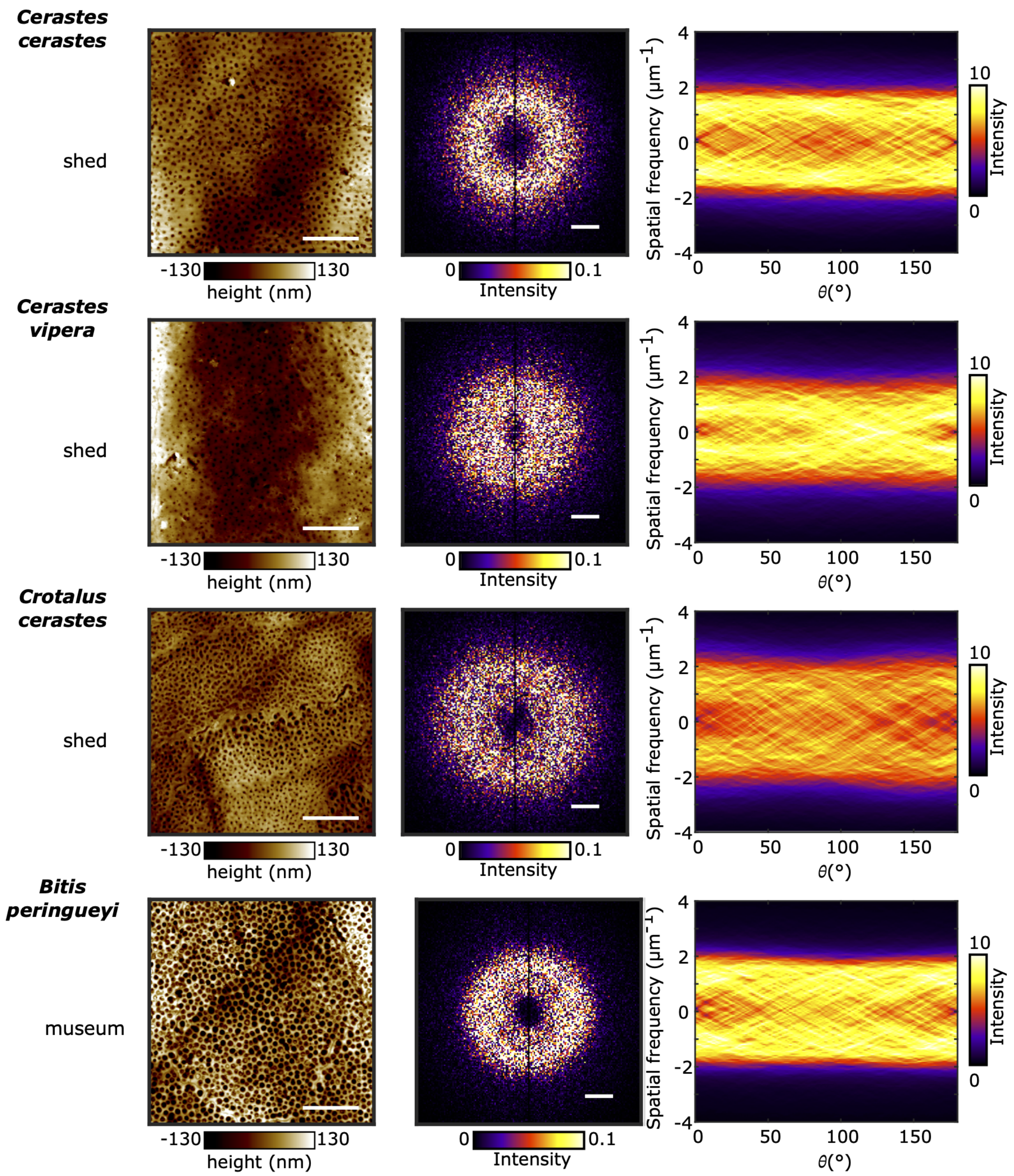


**Figure S12:** Images (20$\mu$m $\times$20$\mu$m), power spectra, and Radon transforms from sidewinding species *Cerastes cerastes*, *Cerastes vipera*, *Crotalus cerastes*, and *Bitis peringueyi* samples. Scale bars: AFM image (left column): 5$\mu$m, power spectrum (middle column): 1$\mu$m$^{-1}$

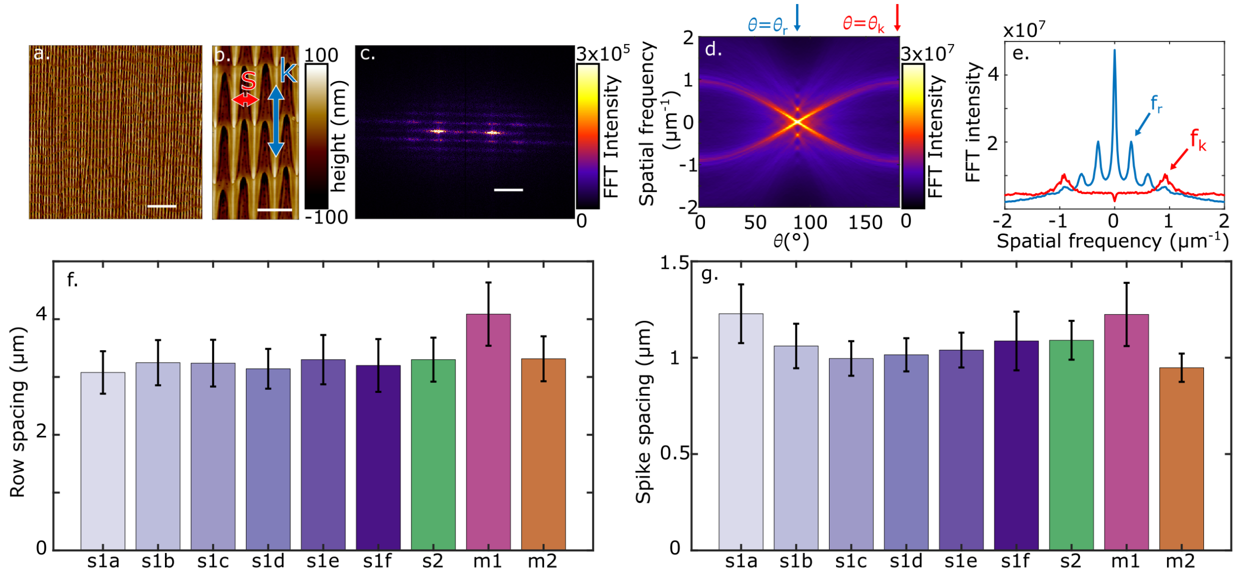


**Figure S13:** The same analysis in Figures 3 and 4 but using a fast Fourier transform (FFT) rather than a power spectrum. **(a)** a 60x60 image of *P. guttatus* microstrucutre. **(b)** a close-up showing the row spacing and spike spacing. **(c)** an FFT of the 60x60 image. **(d)** a Radon transform of the FFT and slices associated with each desired spacing. **(d)** the slices plotted showing the peaks used to find the row and spike spacing. **(e) and (f)** the row and spike spacing of each sample calculated similarly as when a power spectrum is used. Scale bars: **(a)** 10$\mu$m, **(b)** 2$\mu$m, **(c)** 1$\mu$m$^{-1}$,
